# Depletion of astrocyte inflammatory pathway in the arcuate nucleus of the hypothalamus is sufficient to prevent the diet-induced metabolic alterations of polygenically predisposed obese rats

**DOI:** 10.64898/2026.03.27.714441

**Authors:** Anais Bouchat, Luca Papini, Janine Schläpfer, Patricia Kulka, Christelle Le Foll

**Author notes:** Corresponding author: Christelle Le Foll, Institute of Veterinary Physiology, University of Zurich, Winterthurerstrasse 260, 8057 Zurich Switzerland.

## Abstract

Selectively bred diet-induced obesity-prone (DIO-P) rats have defective nutrient sensing prior to obesity onset. We hypothesized that glial inflammation in the arcuate nucleus (ARC) impairs hypothalamic responses to dietary clues, thereby promoting obesity development in genetically susceptible animals. This study established a timeline of inflammatory events in male and female DIO-P and diet-resistant (DR) rats fed either a low fat chow or exposed to a high energy diet (HED; 32% fat, 25% sucrose) for three days or four weeks. On chow diet, DIO-P rats of both sexes displayed elevated astrocyte density and increased expression of pro-inflammatory markers in the ARC, alongside reduced microglial content, compared to DR rats. Three days of HED transiently amplified most MBH pro-inflammatory markers in DIO-P rats. Four weeks of HED decreased GFAP expression in DIO-P rats while Iba1 density remained unchanged, whereas, DR rats showed a reduction in Iba1with no change in GFAP or cytokine expression. To determine whether mediobasal hypothalamus (MBH) astrocyte inflammation contributes to the development and maintenance of an obesity, astrocytic IKKβ was depleted before or after HED exposure. Prophylactic MBH astrocyte-specific IKKβ knockdown prevented subsequent body weight gain, improved glucose tolerance and decreased leptin levels in DIO-P rats to levels comparable to DR rats, with no effect in the latter. In contrast, MBH IKKβ astrocytic depletion in already obese DIO-P rats had no effect on energy homeostasis. Together, these findings validate the DIO-P rat as a polygenic model of obesity predisposition and demonstrate that preventing ARC astrogliosis is sufficient to HED-induced body weight gain and obesity development in genetically susceptible animals, highlighting MBH inflammation as a marker and driver of obesity predisposition.

**Highlights:**

- Chow-fed DIO-P rats present heightened ARC astrogliosis and cytokine expression preceding HED-induced obesity.
- Inhibition of IKKβ in MBH astrocytes prevents DIO-P rats from becoming obese.
- Once obese, inhibition of IKKβ in MBH astrocytes is not sufficient to reverse the obese phenotype.

## 1. Introduction

Key brain regions responsible for maintaining body weight and energy homeostasis are located in the hypothalamus [1; 2] and the caudal hindbrain [3–5]. The mediobasal hypothalamus (MBH), formed by the ventromedial nucleus (VMH) and the arcuate nucleus (ARC), contains neurons that can sense and detect fluctuations in a broad range of metabolic signals [6; 7]. These cues include nutrients (e.g., fatty acids (FA) and glucose) [8–14], hormones (e.g., glucagon-like peptide-1, ghrelin, insulin and leptin), and other metabolic-related signaling molecules, such as ketone bodies [15–21]. Thus, by monitoring nutrient availability and the body’s energy state, MBH neurons adjust their activity to modulate metabolic rate and food intake, thereby maintaining energy homeostasis [7].

Energy balance central control depends not only on neurons but also on glial cells, particularly microglia and astrocytes [17; 22–27]. In addition to providing energy and structural support to neighboring neurons [28], ARC astrocytes are strategically located near blood vessels. They form tight connections with ARC neurons and endothelial cells, enabling neurovascular coupling and the modulation of nutrient uptake and metabolism [25; 29; 30]. Microglia, the resident immune cells of the central nervous system, can adopt a broad spectrum of activation states, from pro- to anti-inflammatory, and also engage in other processes such as synaptic pruning [31; 32]

In response to prolonged high-fat diet (HFD) intake, hypothalamic astrocytes and microglia undergo activation and exhibit a hypertrophic and reactive phenotype known as gliosis [30; 33–35]. This process is notably characterized by upregulated expression of astrocytic and microglial markers: glial fibrillary acidic protein (GFAP) and ionized calcium-binding adapter molecule 1 (Iba1), respectively [23; 36; 37]. Similar to rodents, gliosis has also been observed in the hypothalamus of obese patients using MRI imaging [23] highlighting the clinical translation of obesity-induced inflammation.

This gliosis state is associated with activation of glial inflammatory pathways, promoting secretion of cytokines such as interleukin-6 (IL-6), interleukin-1β (IL-1β), chemokine (C-C motif) ligand 2 (CCL2), interleukin-10 (IL-10) and tumor necrosis factor-α (TNF-α) [2; 22; 23; 38–40]. These cytokines can disrupt insulin and leptin signaling in the VMH [41–44]. The NF-κβ/ IKKβ signaling pathway is a key mediator of these processes. IKKβ, by preventing the inhibitory bond between IκB and NFκB, enables the nuclear translocation of NFκB to promote cytokine expression [1; 2; 45]. Upregulation of neuronal NF-κβ/IKKβ in the MBH of mice worsened the obese phenotype produced by 60% HFD intake [44]. Whole-brain inhibition of astrocyte NF-κβ/IKKβ pathway in mice through IκB overexpression prevented acute hypothalamic astrocyte activation induced by 60% HFD exposure but resulted in a transient 15% increase in caloric intake in the first 24 hours of HFD feeding [34]. Whole brain depletion of IKKβ in mice astrocytes after six weeks of 60% HFD feeding prevented the further development of HFD-induced astrogliosis, excessive body weight gain and glucose intolerance [46].

To establish a causative relationship between hypothalamic astrogliosis and obesity, we utilized diet-induced obesity-prone (DIO-P) and diet-resistant (DR) rats as a model of human obesity [9; 47–49]. These rats are selectively bred to develop polygenically-inherited diet-induced obesity or to remain lean (“diet-resistant”) when challenged with 45% HFD or high-energy diet (HED; 32% fat and 25% sucrose, as a percentage of total energy content) [50]. Unlike outbred rats or mice, selective breeding allows separation of DR and DIO-P rats before chronic HFD/HED exposure, enabling identification of key metabolic alterations that predispose to the development of obesity. Furthermore, DR rats serve as a dietary controls since they consume the same diet but do not become obese [49].

We hypothesized that hypothalamic inflammation associated with cytokine overproduction in DIO-P rats predisposes them for obesity development. The current studies aimed to first establish a timeline of the metabolic and inflammatory events under acute and chronic HED intake that alter astrocyte metabolism and morphology. We then directly assessed the importance of MBH astrocytic inflammation by disrupting the IKKβ pathway in MBH astrocytes of weanling and adult DR and DIO-P rats, determining whether this intervention could prevent obesity in predisposed animals or reverse it once established.

## 2. Material and methods

### 2.1. Animals and housing

Rats were selectively bred to express DIO-P and DR phenotypes. These rats are generated from outbred Sprague-Dawley rats (Janvier Elevage, France) and were selectively bred in our facility following the breeding scheme established by Prof. Barry Levin [50]. The F4 generation was used for Experiment 1, F6 for Experiment 2 and F9 for Experiment 3.

Rats were housed in T2000 macrolon or IVC cages in an environmentally controlled room (21°C, twelve hours inverted light cycle). Food and water were provided *ad libitum*. Rats were fed either a standard chow diet (Diet 3436 or 3336, Provimi Kliba AG, Kaiseraugst, Switzerland; 3.42 kcal/g, 65.4%, 12.3% and 22.4% of calories provided by carbohydrates, fat and proteins respectively) or HED (Diet D12266B, Research Diets, Inc., New Brunswick, USA; 4.41 kcal/g, 16.7%, 32% and 51.4% of calories provided by protein, fat and carbohydrates, half of which is from sucrose, respectively). Handling, measurement of body weight and food intake were conducted weekly before dark onset. All procedures were approved by the Veterinary Office of the Canton Zurich, Switzerland and comply with the ARRIVE guidelines.

### 2.2. Experiment 1: Effect of HED on MBH gliosis in DR and DIO-P rats

Eleven week-old DR and DIO-P rats (n=13 males and 13 females per phenotype) were either kept on chow or switched to HED for three days. Food intake and body weight were measured daily. A second cohort of eight-week old DR and DIO-P rats (n=20 males and 20 females per phenotype) was divided in two: one half was maintained on chow and the other half was switched to HED for four weeks, throughout which body weight and food intake were measured weekly.

In both cohorts, half the animals were sacrificed for qPCR analysis, while the other half was perfused for immunohistochemical staining. The groups were evenly mixed for phenotype, diet and sex.

### 2.3. Experiment 2 (Fig. 4A): MBH IKKβ astrocyte depletion in weanling DR and DIO-P male rats followed by HED

#### 2.3.1. Stereotaxic virus injection in the mediobasal hypothalamus

Three week-old DR and DIO-P males (n=18/phenotype) were bilaterally injected in the MBH with a control AAV shRNA (ssAAV-9/2-hGFAP-hHBbI/E-chI[4x(shm/rNS)]-EGFP-WPRE-bGHp(A) scrambled; v531-9 Repository of the Viral Vector Core UZH) or with an AAV GFAP-IKKβ shRNA virus (ssAAV-9/2-hGFAP-hHBbI/E-chI[4xsh(rIKKb)]-EGFP-WPRE-bGHp(A) custom made by the Viral Vector Core UZH) (n=9/group). The latter is composed of 4 shRNA sequences (5’-3’: 1-cgcagaacttggcacccaatga; 2- aggagatacttgaaccagttcg; 3- cggatgacctagaggaacaagc; 4- aggaacagtgaagttctcaagt). They induced expression of scrambled shRNA or IKKβ translation-inhibiting shRNA respectively, under the control of GFAP promoter. Under injectable anesthesia (ketamine 40 mg/kg and medetomidine, 0.8 mg/kg, s.c.), rats received bilateral viral injections (2.0 x 10E12 vg/ml in 500 nL per side) in the MBH (33G needle; 2.6 mm posterior, ± 0.3 mm lateral to bregma and 8.6 mm ventral to the dura mater) at a constant rate for five minutes. The needle was left in place for five additional minutes to avoid reflux. After reversion of anesthesia (Atipamezole, 0.7 mg/kg, s.c.), the rats were allowed to recover in a warmed cage (27°C) and received meloxicam (2 mg/kg, s.c.) for five days, until fully recovered. After surgery, rats were single-housed and maintained on chow for one week prior being switched at five-week-old to HED for eight weeks. Terminally, the injection site were verified for all rats and five rats were excluded from all analyses.

#### 2.3.2. Assessment of food intake and meal patterns

At eight-week-old (three weeks post-HED onset), rats were individually housed in cages equipped with BioDAQ food intake monitors (Research Diet, New Brunswick, NJ, USA) to continuously measure food intake and meal patterns. Following seven days of acclimation in the BioDAQ cages, *ad libitum* measurements were taken and averaged over three consecutive days. For the fasting-refeeding paradigm, rats were fasted for twelve hours during the light phase, and food was returned at dark onset, meal patterns were then measured for the next 24 hours. Meal pattern criteria were an intermeal interval of 900 s and a minimal meal size of 0.23 g [51].

#### 2.3.3. Indirect calorimetry

At eleven-week-old (six weeks post HED onset), four rats of each group were placed in TSE PhenoMaster indirect calorimetry cages (TSE Systems, Bad Homburg, Germany). O_2_ consumption (VO_2_) and CO_2_ output (VCO_2_) were measured, and used to calculate energy expenditure (EE) and respiratory exchange ratio (RER) based on equations from Weir [52]. To account for differences in body weight and body mass composition, EE data were corrected for individual lean body mass (LBM in g) and fat mass (FM in g) with the following formula: LBM + 0.2FM, as recommended by Even and Nadkarni [53].

#### 2.3.4. Oral glucose tolerance test (OGTT)

Rats were fasted two hours prior the test. At dark onset, they were gavaged with 2 g/kg of D-glucose (0.5 g/ml). Glycemia was sampled from tail blood prior to gavage (0) and at 15, 30, 60, 90, and 120 min post-gavage (Glucometer: Breeze2, Bayer, Zurich, Switzerland).

#### 2.3.5. Body composition

Postmortem body composition (total fat and lean masses) was measured with an EchoMRI (Echo Medical System, Houston, Texas, USA). Scanning was repeated two times per animal.

#### 2.3.6. Leptin and insulin assay

Leptin and insulin plasma levels were measured using a U-PLEX^®^ Metabolic 2-plex combo 1 according to the manufacturer’s protocol (Meso Scale Diagnostics, Rockville, MD, USA).

### 2.4. Experiment 3 (Fig. 7A): MBH IKKβ astrocyte depletion in adult DR and DIO-P male rats fed beforehand with HED

Four-week old DR and DIO-P rats (n=16/phenotype) were fed HED for four to five weeks prior AAV injection at eight-nine week-old. The surgical procedure followed the same protocol as Experiment 2, with the following modifications: an injection volume of 1000 nL per side was used, and the stereotaxic coordinates were -3.5 mm posterior, ±0.4 mm lateral to bregma and -9.7 mm ventral to the skull surface. After surgery, rats were maintained group-housed on HED until the completion of the experiment at 15 week-old. Body weight and food intake was monitored throughout.

### 2.5. Brain collection for immunohistochemistry

After a two hour-long fast at the end of the light phase, rats were injected at dark onset with pentobarbital (100 mg/kg, i.p.; Kantonsapotheke Zurich, Switzerland) to achieve deep anesthesia. Rats were then transcardially perfused with ice-cold 0.1M phosphate buffer followed by 4% paraformaldehyde (PFA). Brains were then post-fixed for 24 hours in 4% PFA followed by 48 hours incubation in 20% sucrose, prior being frozen in hexane and stored at -80°C.

20 μm thick slide-mounted sections of the ARC were cut and collected with a cryostat. Brain sections were rinsed in phosphate-buffered saline (PBS) and blocked for two hours in PBS with 0.3% Triton X, 1% bovine serum albumin (BSA) and 2% normal donkey serum (NDS) at room temperature, before being incubated for 48 hours at 4°C in the same solution complemented with goat anti-GFAP (1:500; ab53554, Abcam, Netherland), rabbit anti-iba1 (1:1000; 019-19741, Wako, IGZ Instruments AG, Switzerland), or mouse anti-HuCD (1:500, A21271, Thermo Fisher Scientific) antibodies. After washing with PBS, brain sections were incubated in PBS, 0.3% Triton X and 2% NDS with Cy3-conjugated donkey anti-goat (1:100; 705-165-147, Jackson ImmunoResearch Europe), Cy3-conjugated goat anti-rabbit (1:100; 111-165-144), AF488-conjugated Donkey Anti Rabbit (1:100, 711-545-152) or AF647-conjugated goat anti-mouse (1:300, A-21235, Thermo Fisher Scientific) antibodies. After rinsing, brain sections were counterstained in DAPI (0.25 μg/ml; 62248, Thermo Fisher Scientific), washed again and coverslipped with Vectashield (H-1400, Vectorlabs, Burlingame, CA, USA).

Immunostained sections were visualized with an Axio Imager 2 microscope (Carl Zeiss, Feldbach, Switzerland) using a 20x objective. Density per area of GFAP and Iba1 was quantified by ImageJ particle analysis (Fiji) by an experimentally-blinded observer.

### 2.6. Sample processing for quantitative PCR

Rats were fasted for two hours before dark onset and were briefly anesthetized with isoflurane followed by decapitation. Brains were collected, immediately frozen on dry ice and stored at -80°C until processing. Brains were mounted on a cryostat and each side of the MBH was then punched, collected and stored at -80°C. MBH mRNA was extracted using a Promega RNA extraction kit (Z6111, ReliaPrep RNA Tissue Miniprep System, Promega, Madison, WI, USA), and mRNA samples were diluted with nuclease free water to achieve a final concentration of 0.125 μg/μl. cDNA was synthetized (BIO-65054, SensiFAST, Bioline, Memphis, TN, USA) and quantitative PCR (qPCR) was performed (7500 Fast Real-Time PCR System). Samples were run in duplicate and gene mRNA expression was measured using the 2^-ΔΔCT^ method using GAPDH as a housekeeping gene. Gene list: GAPDH (Rn01775763_g1), NPY (Rn01410145_m1), POMC (Rn00595020_m1), IKKβ (Rn00584379_m1), UCP2 (Rn01754856_m1), CCL2 (Rn00580555_m1), TNF-α (Rn01525859_g1), IL-1β (Rn00580432_m1), IL-6 (Rn01410330_m1), GFAP (Rn01253033_ m1), Iba1 (Rn00574125_g1).

### 2.7. Statistics

Results were analyzed with Prism 10 using a two-way or a three-way Analysis of Variance (ANOVA) followed by multiple comparisons intergroup analysis as indicated in the figure legends. All data are expressed as mean ± SEM and any statistically significant differences are indicated by asterisks or letters.

## 3. Results

### 3.1. Despite similar short-term overeating responses, only DIO-P rats remain hyperphagic during prolonged HED exposure

To determine the role of obesity predisposition in the establishment of HED-induced inflammatory response, DIO-P and DR rats were exposed to three days or four weeks of HED, while respective control groups were maintained on chow diet.

Over four weeks of chow diet feeding, DIO-P male rats gained 40% more weight and consumed 10% more calories than their DR counterparts (Fig. 1E-G). The same trends were observed in female rats with DIO-P gaining 46% more weight and eating 17% more calories than DR females. (Fig. 1L-N). Nevertheless, despite presenting a higher body weight on chow, the comparable leptin levels in both phenotypes suggests that DIO-P rats are heavier but not fatter prior to HED exposure (Fig. 1C, J).

**Figure 1:**
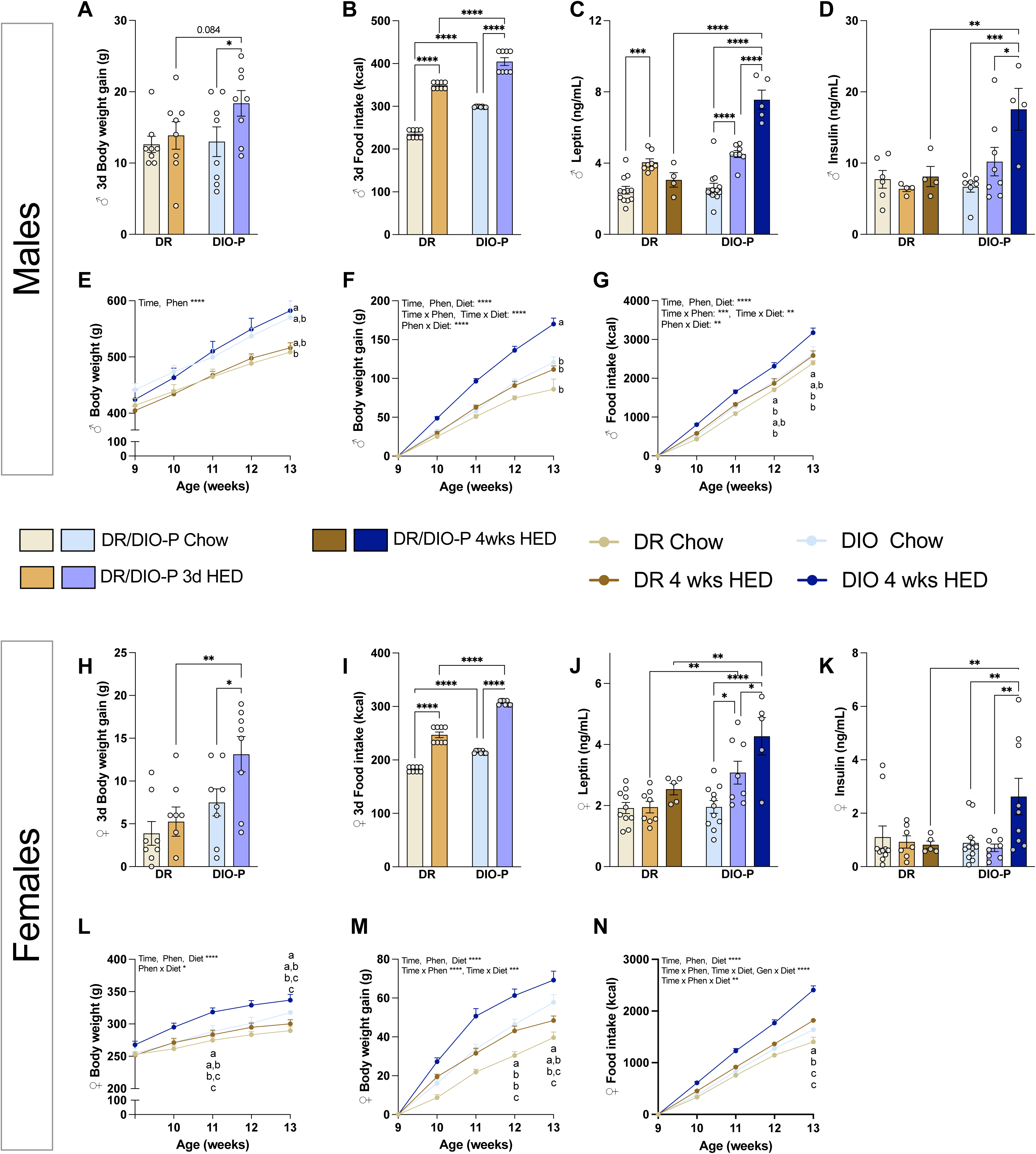
Body weight and energy intake in male and female DR and DIO-P rats during chow, short-term, and prolonged HED exposure. Morphometric data in male and female rats after three days of HED: (**A, H**) body weight gain, (**B, I**) total food intake, (**C, J**) plasma leptin levels, (**D, K**) plasma insulin levels. Data in male and female rats after four weeks of HED: (**E, L**) body weight, (**F, M**) body weight gain, (**G, N**) longitudinal cumulative food intake. Data are represented as mean ± SEM (n= 8-10/group) and were analyzed using a 2-way or 3-way ANOVA (Factors: time, phenotype and diet) followed by Sidak’s and Tukey’s post-hoc test as appropriate. \**p*<0.05, ** *p*<0.01, \*\*\**p*<0.001, \*\*\*\**p*<0.0001. ^a, b, c^Data points with differing superscripts differ from each other at *p*<0.05.

Consistent with previous findings [47], both DIO-P and DR rats increased their caloric intake during the first three days of HED exposure (Fig. 1B, I), but only DIO-P rats tended to gain more weight compared to the chow-fed groups (Fig. 1A, H). DIO-P rats sustained this hyperphagia over four weeks, whereas DR rats normalized their intake and body weight to levels comparable to those of chow-fed groups (Fig. 1G, N). This elevated food intake over prolonged exposure was accompanied by excessive body weight gain in DIO-P males and females compared to HED-fed DR rats and chow-fed DIO-P rats (Fig. 1F, M). Finally, following four weeks of HED, only DIO-P rats displayed hyperinsulinemia and hyperleptinemia, suggesting impaired glucose homeostasis and increased fat mass (Fig. 1C, D, J, K).

Expression of food intake control genes in the MBH revealed that DIO-P rats exhibited a higher baseline expression of the orexigenic peptide NPY, which was reduced by half following HED exposure (Fig. 2H, P). In chow-fed DIO-P rats, anorectic POMC expression was surprisingly seven and eleven times greater than male and female DR rats, respectively. In both sexes, these levels underwent a five-fold reduction following prolonged challenge (Fig. 2G, O). HED had no effect on DR’s gene expression of feeding control genes.

**Figure 2:**
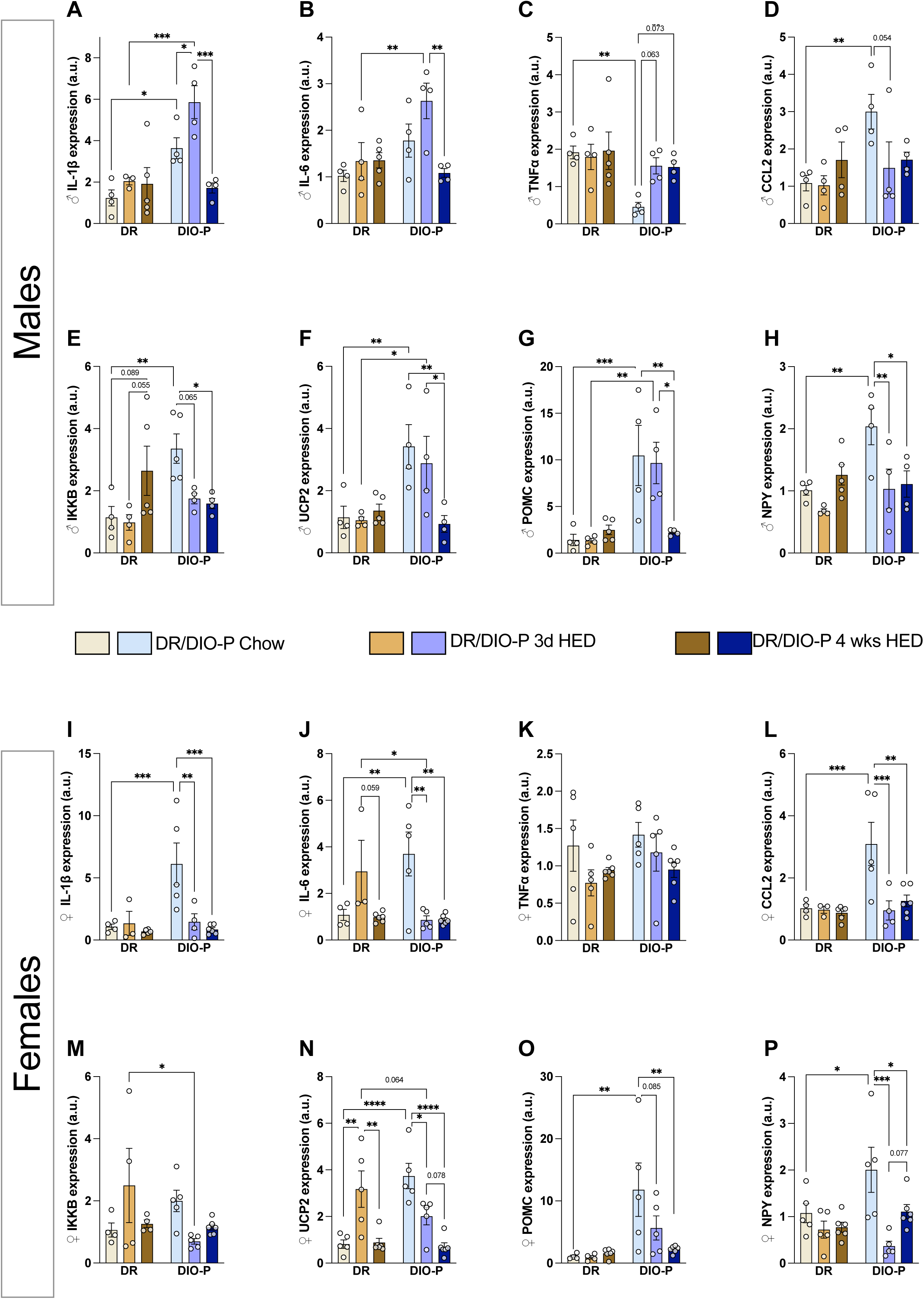
MBH inflammatory and neuropeptide gene expression in male and female DR and DIO-P rats under chow and HED. mRNA expression of (**A, I**) IL-1β, (**B, J**) IL-6, (**C, K**) TNF-α, (**D, L**) CCL2, (**E, M**) IKKβ, (**F, N**) UCP2, (**G, O**) POMC and (**H, P**) NPY, was quantified in the MBH of male (**A–H**) and female (**I–P**) DR and DIO-P rats. Rats were studied under chow, after 3 days or 4 weeks of HED. DR chow-fed rats were used as the reference sample. Data are presented as mean ± SEM (n= 4-6/group) and were analyzed using a 2-way ANOVA (Factors: phenotype and diet) followed by Tukey’s post-hoc test. \**p*<0.05, \*\**p*<0.01, \*\*\**p*<0.001, \*\*\*\**p*<0.0001.

### 3.2. Chow-fed DIO-P rats present heightened inflammatory markers expression in the MBH compared to DR and HED-fed DIO-P rats

Expression of inflammatory genes in the MBH was assessed to determine how they were impacted by three days or four weeks of HED intake. No significant variations were detected in DR rats, apart from a trend towards increased IKKβ expression in DR males undergoing prolonged exposure (+130%, *p*<0.089; Fig. 2E). For both sexes, chow-fed DIO-P rats displayed elevated baseline levels of most inflammatory markers (IL-1β, IL-6, CCL2, and IKKβ) relative to their DR counterparts, with the exception of TNF-α, which was four times lower in DIO-P males (Fig. 2A-E,I-M). In DIO-P males, all aforementioned cytokines were acutely increased by three days of HED, except for the monocyte chemoattractant CCL2 that followed the opposite trend. While IL-6 and IL-1β levels returned to baseline following extended exposure, TNF-α remained elevated. In DIO-P female rats, short-term challenge elicited a down-regulation of all inflammatory markers except TNF-α, suggesting counter-regulatory mechanisms or interactions with estrogen [54]. This decrease was maintained after four weeks of HED exposure. UCP2, an uncoupling protein/transport carrier protein, has been described to lower mitochondrial membrane potential, to shift substrate utilization and to limit mitochondrial reactive oxygen species production in neurons [55; 56]. As observed with the cytokines, UCP2 expression is elevated in chow-fed DIO-P rats and decreased after four weeks of HED (Fig. 2F, N). However, three days of HED increased UCP2 mRNA in DR female rats.

### 3.3. Chow-fed DIO-P rats have higher astrocytic and lower microglia ARC content

Prolonged HFD consumption has been previously associated with gliosis of astrocytes and microglia in the ARC [30; 35]. To determine whether this parameter was altered in DIO-P and DR rats, ARC sections were immunostained to label astrocytes and microglia using antibodies against GFAP and Iba1 respectively. On a chow diet, DIO-P groups displayed an increased astrogliosis compared to their DR counterparts (Fig. 3A, C). These levels were, however, reduced to DR levels following three days or four weeks of HED. GFAP intensity in DR rats was not altered by HED, except for a trend towards a decrease in females following long-term exposure compared to the three-day group (-45%, p<0.083; Fig. 3C). Compared to chow-fed DIO-P rats, DR rats presented a greater Iba1 MBH content at baseline (Fig. 3B,D). Short-term HED challenge led to a transient two-fold increase in DIO-P males, but not in females nor in DR rats. Meanwhile, four weeks on this diet reduced the microglial signal to the same level in all groups.

**Figure 3:**
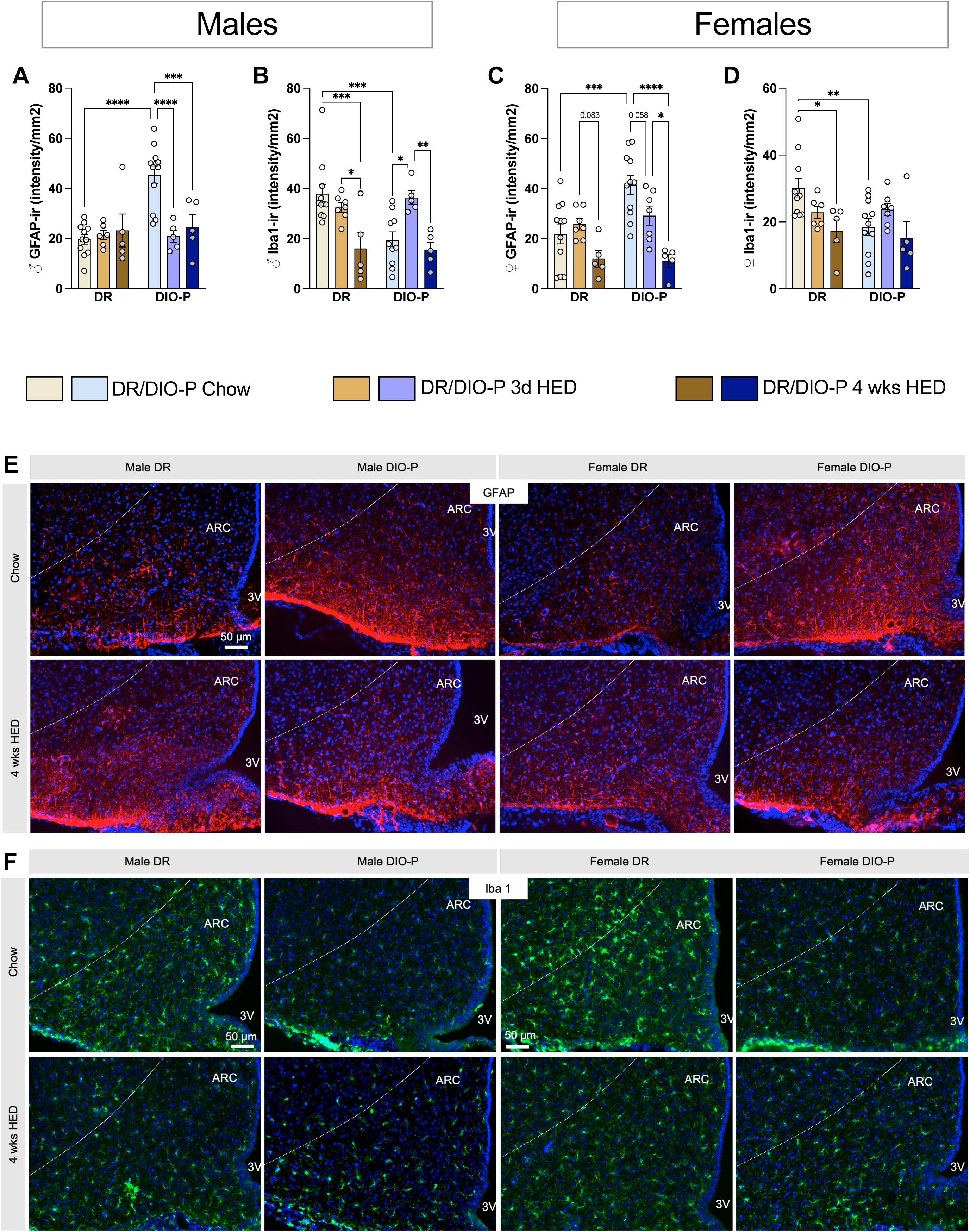
ARC gliosis in female DR and DIO-P rats under chow and HED. Representative quantifications of ARC glial markers are depicted for male and female DR and DIO-P rats across dietary conditions. (**A**) male GFAP density, (**B**) male Iba-1 density,(**C**) female GFAP density, (**D**) female Iba1 density, in rats maintained on chow, after 3 days or 4 weeks of HED. (**E**) Representative immunofluorescent 10X images showing GFAP positive fibers (red), Dapi (blue) in the ARC on chow and after 4 weeks on HED diet. (**F**) Representative immunofluorescent 10X images showing Iba1 positive fibers (green), Dapi (blue) in the ARC on chow and after 4 weeks on HED diet. Data are represented as mean ± SEM (n= 8/group) and were analyzed using a 2-way ANOVA (Factors: phenotype and diet) followed by Tukey’s post-hoc test. \**p*<0.05, \*\**p*<0.01, \*\*\**p*<0.001, \*\*\*\**p*<0.0001.

These results underscore that DIO-P male and female rats fed a chow diet displayed heightened MBH inflammatory gene expression and greater ARC astrocytic density than their DR counterparts. These inflammatory markers were reduced to DR levels following HED consumption, while DR rats were largely unaffected by HED exposure, except for a decrease in ARC Iba1 levels.

### 3.4. Inhibiting IKKβ astrocytic pathway prevents the development of obese phenotype

Given that chow-fed DIO-P rats expressed more GFAP and IKKβ in the ARC than DRs, we hypothesized that this increase in pre-emptive astrogliosis and ARC inflammation might be one of the factors predisposing them to obesity. To test this hypothesis, IKKβ was depleted in GFAP-expressing cells in the MBH of three-to-four week old DR and DIO-P male rats (Fig. 4 A).

**Figure 4:**
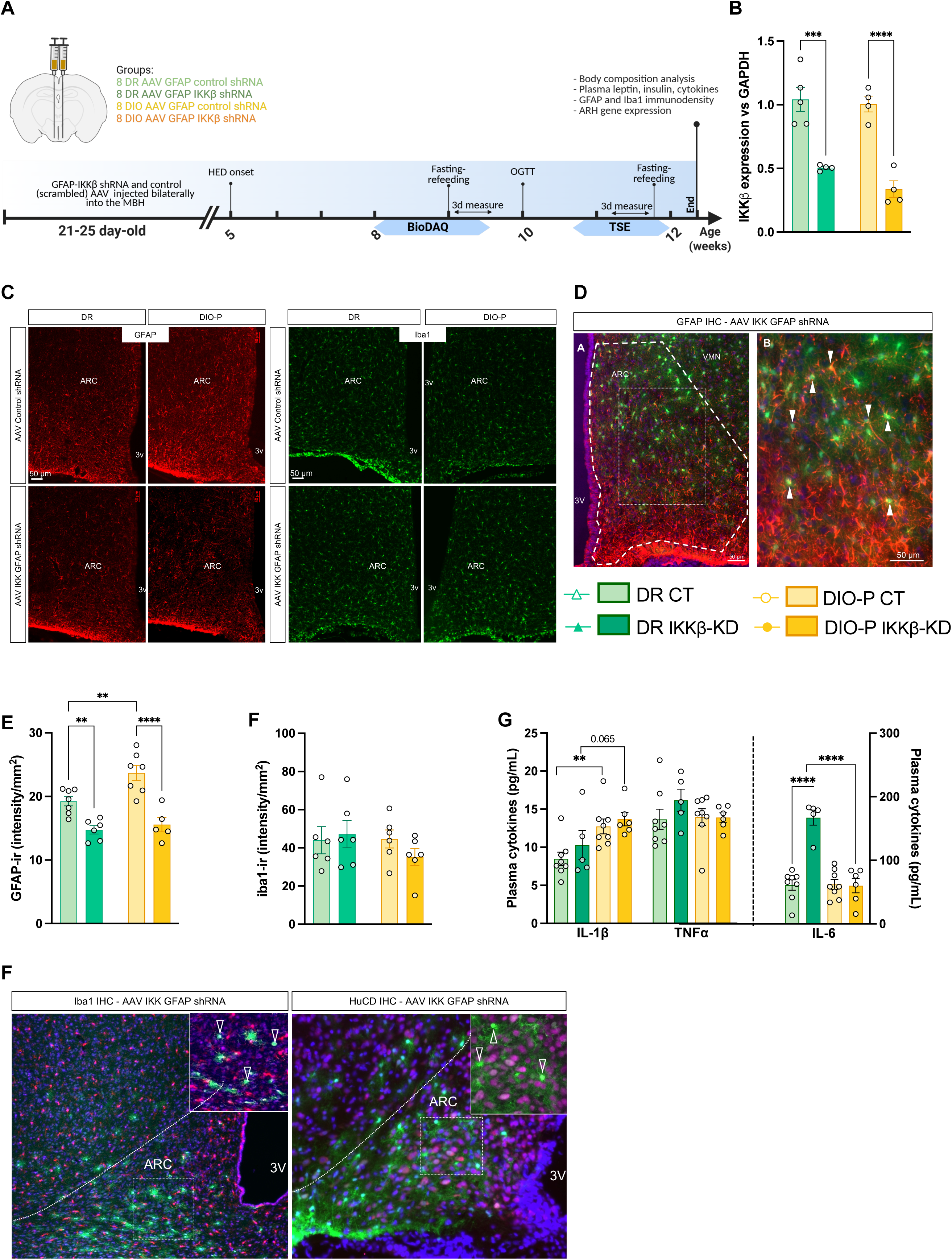
DR and DIO-P male rats injected at weaning in the MBH with an AAV control shRNA or AAV GFAP-IKKβ shRNA and fed HED for 8 weeks. (**A**) Schematic representation of experiment; (**B**) IKKβ mRNA expression in the MBH of DR and DIO-P rats injected in the MBH and sacrificed nine weeks later; (**C**) representative immunofluorescent images showing GFAP (red) and Iba-1 (green) 10X representative images in the ARC of control or knockdown DR and DIO-P rats; (**D**) representative immunofluorescent images showing GFAP positive fibers (red) colocalised with the AAV-GFAP IKKβ shRNA (green) in the ARC of rats, white arrows indicate colocalization; (**E**) quantification of GFAP immunoreactive density in the ARC; (**F**) quantification of Iba1 immunoreactive density in the ARC; (**G**) plasma cytokine concentration at sacrifice. (**H**) representative immunofluorescent images showing Iba1 (red) and HuCD (magenta) positive fibers (red) colocalised with the AAV-GFAP IKKβ shRNA (green) in the ARC of rats, clear arrows indicate absence of colocalization; Data are represented as mean ± SEM (n=6-8/group). Data are analyzed with a 2-way ANOVA (Factors: phenotype and AAV). \**p*<0.05, \*\**p*<0.01, \*\*\**p*<0.001, \*\*\*\**p*<0.0001. ^a, b, c^Data points with differing superscripts differ from each other at *p*<0.05.

#### 3.4.1. Validation of the model and measure of the depletion

Nine weeks after bilateral MBH injections of either the GFAP-IKKβ shRNA AAV (IKKβ-KD group) or the GFAP-control shRNA AAV (CT group) (Fig. 4A), IKKβ expression was quantified by qPCR. The IKKβ shRNA induced a decrease in mRNA transcripts of 52% and 66% in DR and DIO-P rats, respectively (Fig. 4B). Histological analysis of the brains post-sacrifice confirmed that IKKβ depletion was bilateral and encompassed the ARC and VMH areas. GFP colocalized with GFAP but not with HuCD or Iba1 (Fig. 4H), indicating that the AAV was specific to GFAP-positive cells and did not transfect neurons nor microglia.

#### 3.4.2. MBH astrocytic IKKβ knockdown reduces body weight gain and food intake by restoring satiation in DIO-P rats

All groups had a similar body weight at the beginning of the experiment (Fig. 5A). Given that only half of the rats were placed in indirect calorimetry cages at eleven weeks old, which can induce short-term weight loss due to the food hopper habituation, the body weight and food intake are displayed only until this change occurred.

**Figure 5:**
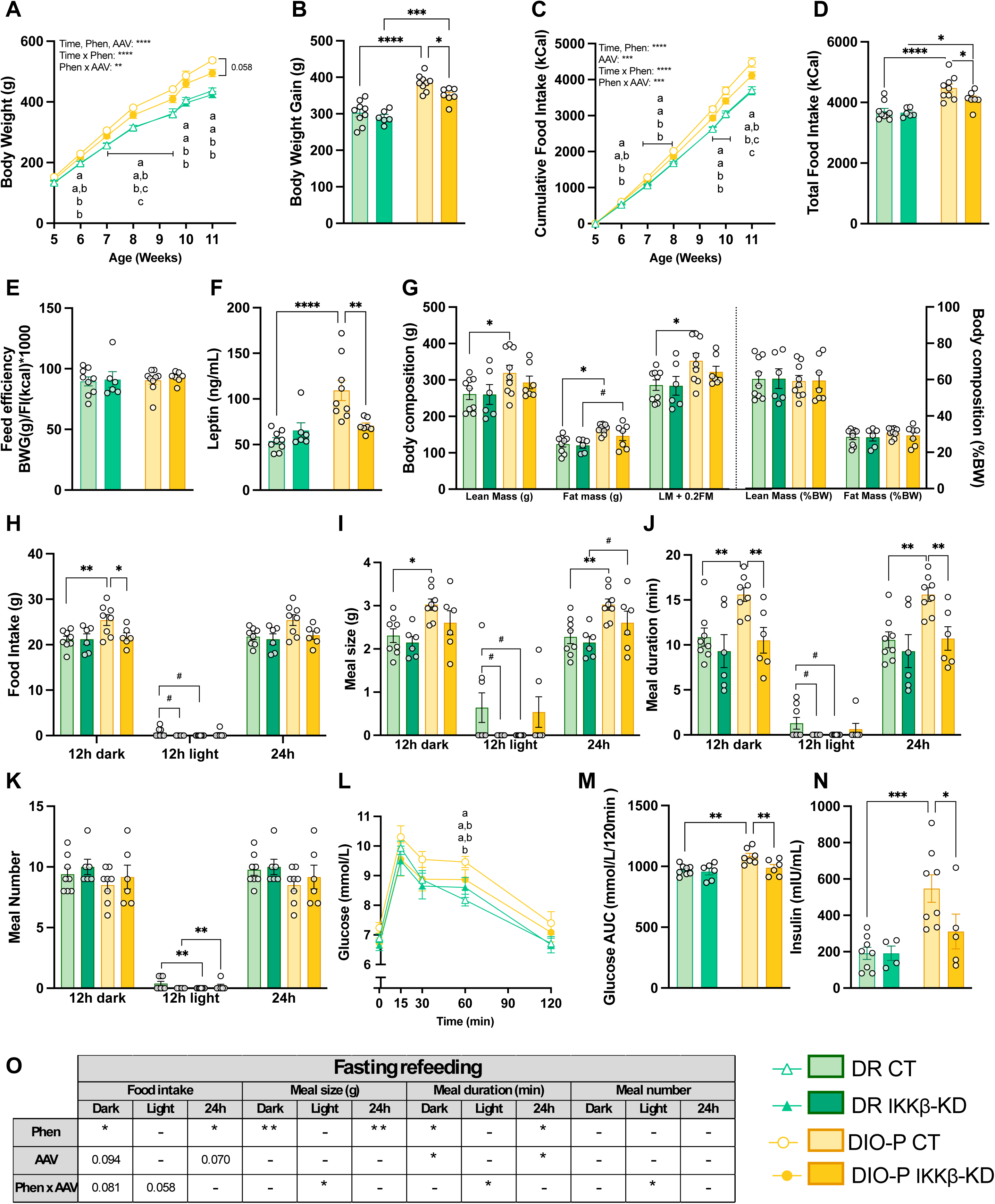
MBH astrocytic IKKβ depletion before HED exposure reduces food intake and alters meal patterning in DIO-P rats. Morphometric and metabolic parameters of DR and DIO-P control (CT) or IKKβ knockdown (IKKβ-KD) AAV in the MBH. (**A**) body weight; (**B**) total body weight gain; (**C**) longitudinal cumulative food intake; (**D**) total food intake; (**E**) feed efficiency calculated from the final body weight gain and total food intake; (**F**) plasma leptin levels at sacrifice; (**G**) body composition: lean and fat mass expressed in (g) or as percentage of body weight (%BW) on the day of sacrifice; (**H-K**) meal pattern analysis after a 12h fast and followed by refeeding: (**H**) cumulative food intake, (**I**) meal size, (**J**) meal duration in minutes and (**K**) meal number over the 12h light phase, 12h dark phase or the entire 24h period. (**L**) plasma glucose concentration over 120 min during an oral glucose tolerance test (2 g/kg), (**M**) area under the curve of the glucose tolerance test, (**N**) plasma insulin level at T0 of the oral glucose tolerance test. (**O**) detailed statistical analysis for figures H-K. Data are represented as mean ± SEM (n=6-8/group) and were analyzed using a 2-way (Factors: phenotype, AAV) or 3-way ANOVA (Factors: phenotype, AAV and time) followed by Tukey’s post-hoc test. \**p*<0.05, \*\**p*<0.01, \*\*\**p*<0.001, \*\*\*\**p*<0.001.

Astrocytic IKKβ knockdown had no effect on either food intake or body weight in DR rats (Fig. 5A-D). At eleven week-old, DIO-P rats were overall 27% heavier than DR rats, but knocking down IKKβ decreased final body weight, body weight gain and food intake in DIO-P rats by 7.8% (*p*<0.058), 8.6% and 11.1% respectively, compared to DIO-P controls (Fig. 5A-D). Neither phenotype nor IKKβ depletion had an effect on feeding efficiency (Fig. 5E). These changes in food intake were accompanied by a trend toward reduced expression of the orexigenic peptides NPY (-80%, *p*<0.070) and AgRP (-47%, *p*<0.080) in DIO-P IKKβ-KD rats compared to DIO-P control (Table 1). Expression of POMC was unaffected by IKKβ depletion in DR while it was significantly decreased by 42% in DIO-P rats (Table 1).

**Table 1:** ARC gene expression of male DIO-P and DR rats injected in the MBH with an AAV control shRNA or AAV IKKβ shRNA and fed HED for eight weeks.

| Fold change of gene of interest vs GAPDH | DR Control | DR AAV GFAP-IKKβ shRNA | DIO-P Control | DIO-P AAV GFAP-IKKβ shRNA | Descriptive statistics |
| --- | --- | --- | --- | --- | --- |
| NPY | 1.03 ± 0.14 | 1.00 ± 0.42 | 1.80 ± 0.61 | 0.33 ± 0.02 | AAV $p=0.06$; F (1, 12) = 4.02<br>AAV x Phen $p=0.07$; F (1, 12) = 3.67 |
| AgRP | 1.02 ± 0.17 | 0.88 ± 0.03 | 0.99 ± 0.20 | 0.52 ± 0.02 | AAV $p<0.05$; F (1, 12) = 6.32 |
| POMC | 1.42 ± 0.59<sup>a</sup> | 3.45 ± 0.18<sup>a</sup> | 6.55 ± 0.84<sup>b</sup> | 3.76 ± 0.82<sup>a</sup> | Phen $p<0.001$; F (1, 12) = 16.95<br>AAV x Phen $p<0.01$; F (1, 12) = 13.31 |
| IKKβ | 1.02 ± 0.12<sup>a</sup> | 0.51 ± 0.00<sup>b</sup> | 1.01 ± 0.06<sup>a</sup> | 0.34 ± 0.08<sup>b</sup> | AAV $p<0.001$; F (1, 12) = 61.63 |
| GFAP | 1.03 ± 0.14<sup>a</sup> | 1.18 ± 0.15<sup>a</sup> | 2.80 ± 0.24<sup>b</sup> | 1.55 ± 0.53<sup>a</sup> | Phen $p<0.01$; F (1, 12) = 11.92<br>AAV x Phen $p<0.05$; F (1, 12) = 5.17 |
| Iba1 | 1.00 ± 0.06<sup>a</sup> | 0.90 ± 0.07<sup>a</sup> | 0.85 ± 0.12<sup>a</sup> | 0.54 ± 0.03<sup>b</sup> | Phen $p<0.01$; F (1, 12) = 11.40<br>AAV $p<0.05$; F (1, 12) = 7.133 |
| IL-6 | 1.06 ± 0.21 | 1.27 ± 0.20 | 0.81 ± 0.18 | 0.79 ± 0.03 | ns |
| IL-1β | 1.16 ± 0.37 | 1.37 ± 0.05 | 1.28 ± 0.18 | 0.74 ± 0.12 | ns |
| TNF-α | 1.30 ± 0.36 | 1.42 ± 0.47 | 1.27 ± 0.27 | 1.06 ± 0.28 | ns |
| CCL2 | 1.13 ± 0.30 | 1.19 ± 0.16 | 0.99 ± 0.12 | 0.67 ± 0.15 | ns |

DR control rats were used as the reference sample. Data are presented as mean ± SEM (n= 5/group) and were analyzed using a 2-way ANOVA (Factors: phenotype and AAV) followed by Tukey's post-hoc test. \* $p<0.05$ , \*\* $p<0.01$ , \*\*\* $p<0.001$ , \*\*\*\* $p<0.0001$ . <sup>a, b, c</sup> Data points with differing superscripts differ from each other at the $p<0.05$ .

To determine whether these changes in food intake were caused by alterations in meal patterns, rats were placed in BioDAQ cages for two weeks. Food intake, meal number, size and duration were assessed during *ad libitum* feeding (Suppl. Fig. 1A-D) and after a twelve-hour fast (Fig. 5H-K). During *ad libitum* feeding, no differences induced by IKKβ depletion were observed in the meal patterns in both DIO-P and DR rats. In fasted and refed rats, reducing IKKβ in GFAP-expressing MBH cells prevented the increase in food intake and meal duration seen in DIO-P controls without altering meal number (Fig. 5H, J, K). DIO-P IKKβ-KD rats also consumed smaller meals than control DIO-P rats, although their meal size over 24 hours remained greater than that of DR rats (Fig. 5I). These results suggest that the decrease in total food intake induced by the depletion of MBH astrocytic inflammation may be mediated by the restoration of DIO-P meal patterns, notably satiation, to those of DR rats.

#### 3.4.3. MBH astrocytic IKKβ-knockdown do not alter body composition but prevented hyperleptinemia in DIO-P rats

As expected after eight weeks on HED, DIO-P rats presented higher absolute fat and lean masses than DR rats. Normalizing these values to body weight eliminated the difference, suggesting that body composition ratios remained similar across all groups (Fig. 5G). Consistent with their increased fat mass, DIO-P controls showed increased plasma leptin levels. However, IKKβ knockdown in DIO-P rats prevented this increase, with plasma leptin levels remaining similar to those of DR rats (Fig. 5F).

#### 3.4.4. IKKβ-knockdown DIO-P rats preserves intact glucose homeostasis

After five weeks of HED, glucose homeostasis was assessed by an OGTT. Despite similar baseline glucose levels across all groups, DIO-P rats displayed lower glucose tolerance than DR rats (Fig. 5L, M). This impairment was partially rescued in IKKβ-knockdown DIO-P rats compared to their control counterparts. Analysis of insulin levels at sacrifice revealed a significant effect of phenotype (*p*<0.01), driven by markedly elevated insulin levels in DIO-P control rats compared to their DR counterparts, while DIO-P IKKβ-KD insulin levels were 43% lower than DIO-P control (Fig. 5N).

#### 3.4.5. Reducing astrocytic inflammation modulates energy expenditure and metabolic fuel preference in DIO-P rats

To further assess the consequences of depleting astrocytic inflammation and potentially explain the reduced body weight gain, we measured the EE and RER. Overall, DIO-P displayed lower EE than DR rats (Fig. 6A, B, Suppl. Fig. 1G, H). In DIO-P rats, IKKβ knockdown tended to increase EE (Fig. 6A, B; Suppl. Fig. 1G, and this effect was even more pronounced during the dark phase of the fasting-refeeding paradigm (+20%, *p*<0.053, Fig. 6A, B). The RER was elevated in DIO-P rats under *ad libitum* conditions, indicating a greater reliance on carbohydrates as an energy source (Supp. Fig. 1E, F). In the fasted state, all groups decreased their RER, reflecting a switch towards fat metabolism, except DIO-P IKKβ-KD, who maintained an RER similar to fed conditions (Fig. 6 C,D).

**Figure 6:**
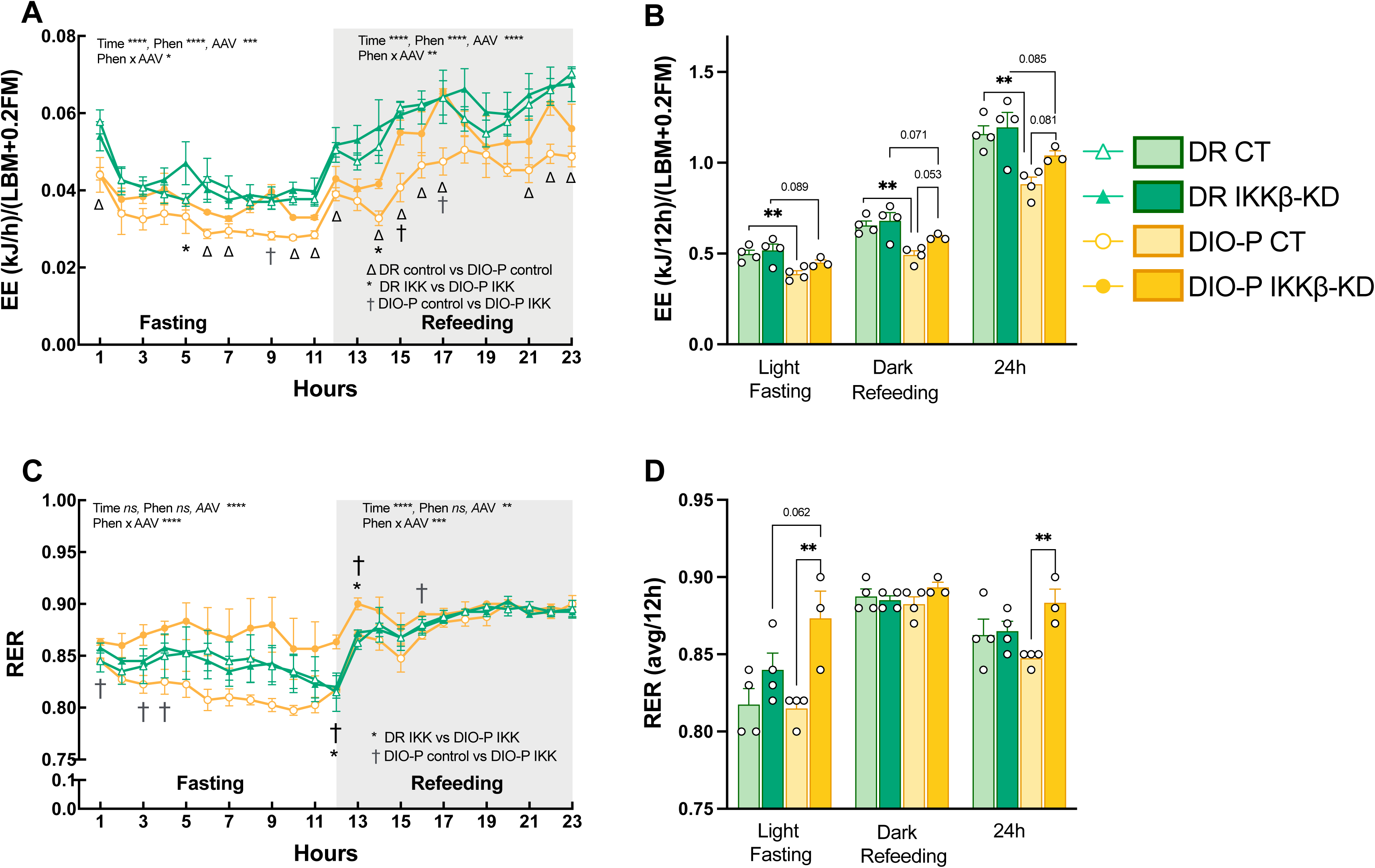
MBH astrocytic IKKβ depletion increases energy expenditure and shifts substrate utilization in DIO-P rats during fasting-refeeding. Time course of energy expenditure during light-phase fasting and dark-phase refeeding in DR and DIO-P rats injected with control (CT) or IKKβ knockdown (IKKβ-KD) AAV in the MBH. (**A**) Measurement of EE (in kJ/h) in hourly bouts over 12h fasting and 12h refeeding periods (grey area) and (**B**) expressed as the cumulative sum of the light, dark or entire 24h period. EE was normalized to body composition (lean body mass+0.2 fat mass) [53]. (**C**) Measurement of RER in hourly bouts over 12h fasting and 12h refeeding periods (grey area) and (**D**) expressed as the average of the light, dark or entire 24h period. Data are represented as mean ± SEM (n=4/group) and were analyzed for each time period using a 2-way (Factors: phenotype, AAV) or 3-way ANOVA (Factors: phenotype, AAV and time) followed by Tukey’s post-hoc test. \**p*<0.05, ** *p*<0.01, *** *p*<0.001. ^Δ^DR control vs DIO-P control, *DR IKK vs DIO-P IKK, ^†^ DIO-P control vs DIO-P IKK, # DR IKK vs DIO-P IKK at *p*<0.05.

#### 3.4.6. Ikkb-KD limits MBH gliosis and modulates inflammatory marker expression

Gliosis was measured at mRNA and protein levels through qPCR and immunofluorescence staining. GFAP levels were higher in DIO-P controls compared to DR controls in both analyses (Table 1, Fig. 4E). Knocking down IKKβ in MBH astrocytes decreased GFAP immunoreactivity by 23% and 34% in DR and DIO-P rats, respectively, while it reduced GFAP mRNA content only in DIO-P rats (Table 1, Fig. 4C, E). The microglia marker Iba1 was decreased at the mRNA level in DIO-P IKKβ-KD (Table 1) while this change was not reflected at protein level, as Iba1 immunoreactivity remained constant across all groups (Fig. 4C, F).

Assessment of MBH cytokines expression revealed no variations in their expression in DR and DIO-P IKKβ-KD groups compared to their respective controls (Table 1). However, plasma IL-6 levels were tripled in DR IKKβ-KD relative to the three other groups (Fig. 4G). IKKβ depletion in MBH astrocytes had no effect on plasma IL-1β or TNF-α across all conditions (Fig. 4G).

### 3.5. Inhibiting MBH IKKβ astrocytic pathway in already obese DIO-P rats does not dampen the obese phenotype

Having shown that knocking down IKKβ in MBH GFAP-positive cells could limit the subsequent development of obesity, we next tested whether the same manipulation could rescue metabolic impairments in already obese DIO-P rats as seen in Fig. 1. Obesity was induced by 4.5 weeks of HED feeding prior to AAV injections, which were followed by 5.5 additional weeks on HED. Overall, IKKβ knockdown had minimal effects in both DIO-P and DR rats, except for a significant interaction between phenotype and AAV in caloric intake, driven by a very mild increase in HED consumption in DR rats following IKKβ knockdown, compared to DR control rats (Fig. 7 C, D). IKKβ inhibition also reduced circulating TNF-α levels, which was significant only in DR rats (Fig. 7J). As expected, DIO-P rats were heavier and ate more than DR rats throughout the whole experiment (Fig. 7A, B). Similarly, body composition was affected only by phenotype, with DIO-P rats displaying increased absolute fat and lean masses, as well as elevated percentage of fat mass relative to body weight (Fig. 7G). Insulin was also significantly higher in DIO-P than DR rats (*p* = 0.0263, +56.9%; Fig. 7H). Finally, leptin and glucose levels were similar across all groups (Fig. 7 F, I). These results suggest that, once the obese phenotype is established in DIO-P rats, it cannot be reversed by IKKβ depletion.

**Figure 7:**
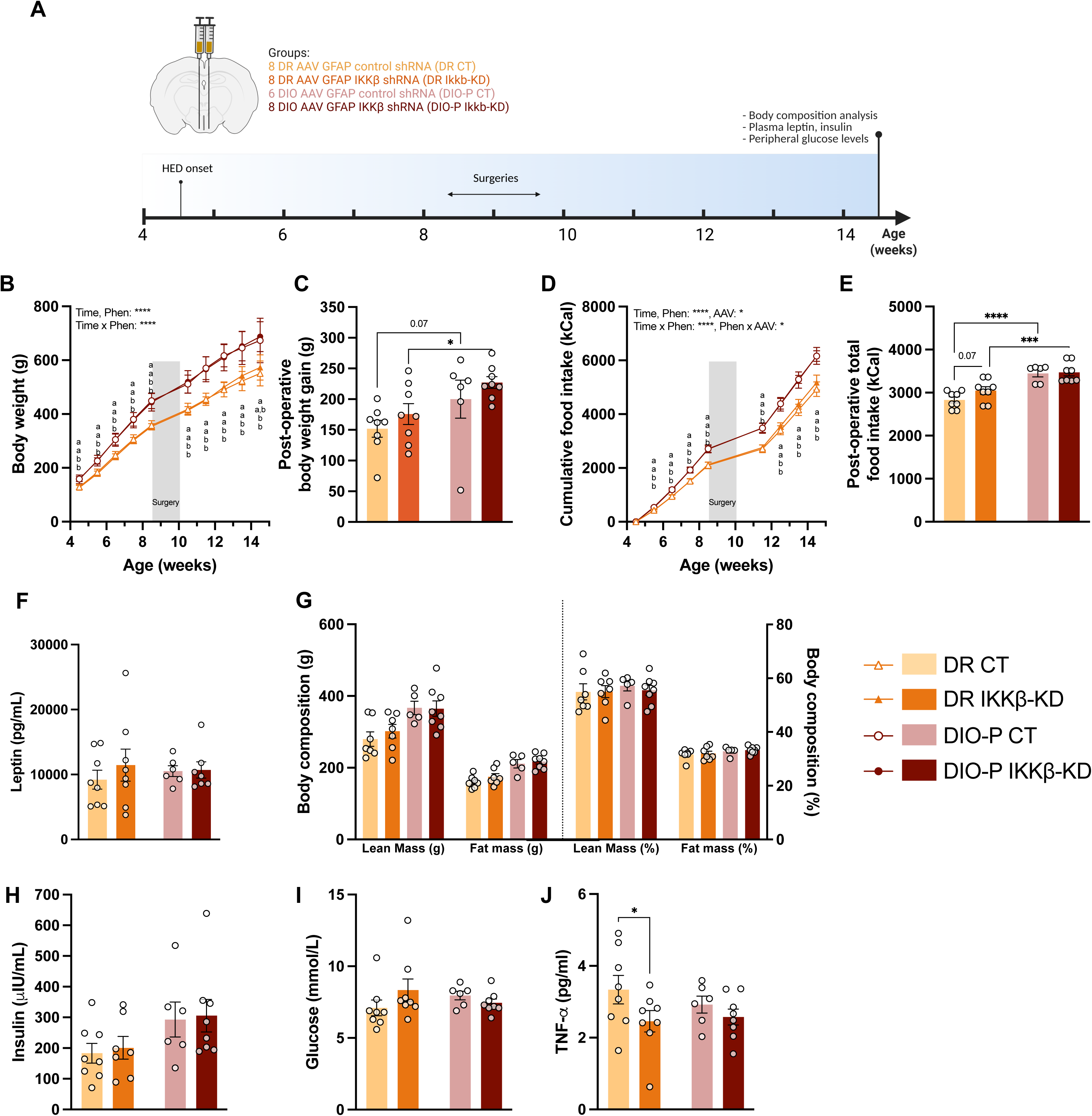
MBH astrocytic IKKβ depletion after 6 weeks of HED exposure does not reverse the obese phenotype. (**A**) Schematic representation of experiment. Morphometric and metabolic parameters of DR and DIO-P control (CT) or IKKβ knockdown (IKKβ-KD) AAV in the MBH. (**B**) body weight; (**C**) total body weight gain; (**D**) cumulative food intake; (**E**) total food intake; (**F**) plasma leptin levels at sacrifice; (**G**) body composition: lean and fat mass expressed in (g) or as percentage of body weight (%BW) on the day of sacrifice; (**H**) plasma insulin at sacrifice; (**I**) plasma glucose at sacrifice; (**J**) plasma TNF-α at sacrifice. Data are represented as mean ± SEM (n=6-8/group) and were analyzed using a 2-way (Factors: phenotype, AAV) or 3-way ANOVA (Factors: phenotype, AAV and time) followed by Tukey’s post-hoc test. \**p*<0.05, \*\**p*<0.01, \*\*\**p*<0.001, \*\*\*\**p*<0.001.

## 4. Discussion

While considerable research efforts have enabled the identification of numerous molecular pathways involved in obesity, the mechanisms conferring predisposition or resistance upon chronic HED or HFD exposure remain poorly understood in both humans and rodent models of obesity. Selectively bred polygenic rodent models such as the DIO-P and DR rats represent some of the most translatable surrogates for human obesity [57], allowing investigation of the metabolic alterations which encode obesity susceptibility prior to obesogenic diet exposure.

The objective of this study was to determine the role of MBH astrogliosis, microgliosis and cytokine production in food intake regulation in DR and DIO-P rats fed HED as previous findings showed that DIO-P, but not DR rats, maintain excessive caloric intake after three days on such diets [9; 47; 48]. We leveraged this model to test the hypothesis that a pre-existing microinflammatory state in the ARC, characterized by elevated astrogliosis and heightened cytokine expression, causally predispose to obesity.

### Preemptive astrogliosis and MBH inflammation in obesity-prone rats

A central finding of this study is that adult DIO-P rats display a distinct glial and inflammatory phenotype in the MBH well before any exposure to an obesogenic diet or any measurable difference in adiposity. On chow, DIO-P rats of both sexes presented elevated GFAP-immunoreactive density, reduced Iba1 density, and upregulated mRNA expression of pro-inflammatory markers relative to DR counterparts. Importantly, this occurred in the absence of increased fat mass or plasma leptin levels, indicating that the neuroinflammatory priming is not a consequence of obesity but a potential pre-condition. Gliosis can even be observed prior to body weight gain in HFD-fed mice [23; 34] and less than 24 hours of exposure is sufficient to trigger the neuroinflammatory cascade [34; 58; 59]. Our findings are also consistent with clinical data showing that healthy overweight individuals have shown elevated hypothalamic water content, a surrogate marker for central gliosis, despite the absence of peripheral metabolic alterations [60], reinforcing the translational relevance of the DIO-P model.

Opposite to DIO-P rats, DR rats displayed an elevated microglial density under chow conditions. The elevated baseline microglial density observed in DR rats may exert anti-inflammatory or modulatory functions that may prevent astrocytes to develop an inflammatory phenotype upon HED exposure [61; 62]. We could thus hypothesize that the reduced microglial presence in DIO-P rats may leave astrocytes in a constitutively reactive state, uncontrolled by microglial anti-inflammatory signaling. This opposite glial structure, with an heightened astrogliosis alongside decreased microglial, may represent a pre-existing microinflammatory environment setting up the stage for diet-induced obesity development.

### Dynamic hypothalamic gliosis during HED challenge

Contrary to the prevailing view that HFD/HED feeding result in increased hypothalamic GFAP immunoreactivity [27; 30; 34; 63], chow-fed DIO-P rats showed a reduction in GFAP density after both three days and four weeks on HED, to levels comparable to DR rats. Notably, this reduction in astrocytic density occurred alongside an acute but transient increase in pro-inflammatory gene expression, suggesting that GFAP downregulation does not necessarily prevent, and may even accompany, hypothalamic inflammation. Reactive astrocytes can adopt functionally distinct states, broadly classified as pro-inflammatory or homeostatic, that are not reliably distinguished by GFAP alone [36]. Recent studies have highlighted the dynamic aspects of astrogliosis and have identified a biphasic response to HFD feeding: a first transient wave, possibly indicative of a high-energy state [58], followed by a second persistent activation phase upon long-term exposure [23; 58; 64]. For example, male mice displayed elevated hypothalamic GFAP density after one week and again after eight months on 60% HFD, but not after three weeks [23]. Similarly, five days on a 58% high-fat high-sucrose (HFHS) diet upregulated astrocytic signaling pathways related to hormone and nutrient sensing, before returning to baseline chow-levels after 15 days [64]. Our studies confirm the dynamic nature of gliosis and that polygenic resistance to obesity appears to confer protection against HED-induced gliosis, whereas DIO-P rats exhibit baseline astrogliosis under chow conditions. Whether astrogliosis may reoccur at later stages of HED in our DIO-P model requires further investigation under prolonged exposure conditions.

### Impact on ARC neurocircuitry and neuropeptide expression

The ARC neurocircuitry is one of the best characterized brain structures involved in the control of energy balance. As part of an integrative system monitoring nutrient availability, ARC POMC and NPY neurons function as nutrient sensors [14; 65–68]. In our study, both POMC and NPY mRNA levels are upregulated in chow-fed DIO-P rats [69], in parallel with elevated inflammatory markers compared to DR rats. After four weeks of HED intake, expression of both neuropeptide was reduced, albeit to different extent, resulting in an increased NPY relative to POMC ratio. This elevated orexigenic tone in HED-fed DIO-P rats may contribute to sustaining hyperphagia. Notably, ARC IKKβ knockdown in HED-fed DIO-P rats reduced both POMC and NPY expression, indicating that their regulation closely related to inflammatory signaling and/or to secondary changes in body weight and leptin levels.

Even under chow diet conditions, ARC circuitry differs between DIO-P and DR rats, with DIO-P rats displaying more inhibitory synapses (orexigenic) than excitatory synapses on POMC neurons [70]. In mice, chemogenetic activation of ARC GFAP-positive cells increases food intake by elevating AgRP/NPY neurons activity [71], highlighting a possible glia-AgRP/NPY connection. Thus, the greater pre-emptive astrogliosis observed in the ARC of DIO-P rats could be involved in this synaptic remodeling and altered neuropeptide expression in nutrient-sensing neurons, thereby increasing the susceptibility to obesity when exposed to an HED or HFD.

### Sex differences in hypothalamic inflammatory response

A notable sex-specific divergence was observed in cytokine dynamics during HED exposure in DIO-P rats. While DIO-P males showed acute upregulation of IL-1β and IL-6 following three days of HED, DIO-P females exhibited a paradoxical down-regulation of the same markers, maintained at four weeks. This divergence likely reflects the modulatory actions of estradiol on hypothalamic NF-κB/IKKβ signaling: estrogen receptor-α can interact with NF-κB subunits to suppress transcriptional activity [54], potentially dampening the acute cytokine response in female rats. The similar GFAP profiles and metabolic outcomes across sexes nevertheless suggest that astrogliosis and its functional consequences are conserved, while cytokine dynamics potentially differ by sex. It this polygenic model, predisposition in females may also be driven by mechanisms different from males, and more specifically, other than inflammation.

### Preventing obesity through MBH astrocyte-specific IKKβ depletion

Given the pre-existing astrogliosis and pro-inflammatory MBH state in chow-fed DIO-P rats, we investigated whether selectively attenuating MBH astrocyte-driven inflammation before HED exposure could prevent obesity development. Targeted depletion of IKKβ in MBH astrocytes at weaning, before any differences in body weight or food intake become evident between DR and DIO-P rats, reduced subsequent HED-induced hyperphagia, body weight gain and improved glucose homeostasis in DIO-P rats to near DR levels. Analysis of meal patterns indicates that the lower weight gain was mediated principally through improved satiation with IKKβ-KD DIO-P rats showing normalized meal size and duration without changes in meal frequency. IKKβ depletion also induced a significant elevation in EE during the dark phase of the fasting-refeeding paradigm, suggesting partial restoration of sympathetic tone or thermogenic activity of the brown adipose tissue [72; 73]. Total fat content showed only a slight decreasing trend, while the obese phenotype was nevertheless prevented, as indicated by the normalization of plasma leptin levels. This apparent discrepancy between fat mass and plasma leptin levels may reflect changes in fat distribution between subcutaneous and visceral fat depots [74; 75].

Previous studies using genetically modified mice to knockdown IKKβ in neurons or astrocytes across the entire brain demonstrated beneficial effects on metabolism and eating behavior but did not identify a specific brain area as being key to the observed improvements [44–46]. Our study, to our knowledge, is the first to identify the contribution of astrocytic inflammatory response in the MBH of polygenically obesity-prone rats.

By decreasing neuroinflammation in DIO-P rats, we may have preserved nutrient neuronal sensing and activity within key hypothalamic circuits. Indeed, ARC astrocytes undergo morphological remodeling in response to metabolic cues and consequently affect neuronal activity and synaptic input strength [76–78]. For example, a single HFD meal increases astrocyte coverage and reduces POMC activity [79], and HFD feeding can promote maladaptive glial ensheathment of ARC neurons [70]. In DIO-P rats, pre-existing astrogliosis may lower the threshold for this maladaptive ensheathment, explaining why these rats fail to adapt to nutrient excess by lowering their intake as DR rats do. Attenuating this inflammatory drive via IKKβ knockdown may therefore improve astrocyte plasticity, enabling normal retraction after meals [58] and proper synaptic balance on ARC neurons, thereby restoring the satiation capacity of the melanocortin circuit. Future studies to examine astrocyte morphology during meal consumption in DIO-P and DR rats would directly test this model.

While MBH astrocytic inflammation seem to be required for the induction of obesity in DIO-P rats, depleting it after obesity establishment could not rescue the metabolic impairments. Indeed, when MBH astrocytic IKKβ knockdown was performed in DIO-P rats that had already developed obesity through five weeks of HED feeding, it had no effect on body weight, food intake, fat mass, insulin, or leptin levels. These results are however scientifically important as they indicate that once the obese phenotype is installed, NF-κB/IKKβ activity in the MBH is no longer a primary driver of metabolic impairments in DIO-P rats. At this stage, additional and potentially irreversible mechanisms are likely taking place such as the synaptic remodeling of hypothalamic circuits [80] including increased inhibitory tone on POMC neurons [70; 76]. Alongside inflammation, obese DIO-P rats present other metabolic alterations, such as hypothalamic leptin resistance, which is a key driver of the obesity phenotype and may not be fully rescued by IKKβ attenuation alone [49; 81; 82]. These results contrast from those obtained in whole-brain astrocytic IKKβ knockout mice, in which the manipulation did not prevent HFD-induced obesity development in lean GFAP-CreERT2 mice, while it prevented glucose intolerance and protected the mice from further weight gain when they were fed HFD for six weeks prior depletion [46]. This discrepancy may reflect the regional specificity of our MBH-targeted approach, whereas the IKKβ-GFAP KO mouse model affects astrocytes throughout the brain [46]. Other areas such as the dorsal vagal complex, which also participate to the regulation of energy balance and glucose homeostasis, may have also contributed to those effects [83; 84]. Additionally, inflammation may cease to be a primary driver of obesity maintenance in obesity-prone rats once obesity is established, while it remains relevant in rodents lacking a specific predisposition. Altogether, these finding suggests that early intervention, before HFD or HED exposure, is critical to prevent the obese phenotype in polygenic susceptible animals.

## Conclusion

Our findings demonstrate that polygenic predisposition to obesity is associated with pre-existing ARC neuroinflammatory state that precedes HED exposure or apparent obesity. Early intervention targeting astrocytes inflammation through MBH-specific astrocyte IKKβ depletion can prevent diet-induced obesity in susceptible animals, but only when implemented before HED exposure. These results highlight the importance of early hypothalamic inflammatory and glial changes as determinants of obesity risk and points to a potential therapeutic windows for intervention. Thus, assessing hypothalamic gliosis in pre-obese individuals, via imaging of the MBH gliosis, which has been associated with glucose intolerance, may prove useful as a predictive tool [60; 85].

## Author Contributions

CLF contributed to conceptualization and funding acquisition. AB, LP, JS, PK and CLF contributed to methodology and investigation. AB, LP and CLF contributed to formal analysis and writing of the original draft. CLF reviewed and edited the final version of this manuscript. All authors approved the submitted version.

## CRediT authorship contribution statement

**Anais Bouchat**: Investigation, data analysis and interpretation, writing-original draft. **Luca Papini**: Investigation, data analysis. **Janine Schläpfer**: Investigation. **Patricia Kulka**: Investigation. **Christelle Le Foll**: Conceptualization, methodology, investigation, writing-original and final draft, editing, guarantor of this work and, as such, had full access to all the data in the study and take responsibility for the integrity of the data and the accuracy of the data analysis. Full datasets are available upon request.

## Acknowledgments and funding

This work was funded by the UZH Forschungskredit (grant FK-019-054 to CLF), the Swiss National Foundation (SNF-310030_207960 to CLF) and the Institute of Veterinary Physiology. The laboratory work was partly performed using the logistics of the Center for Clinical Studies at the Vetsuisse Faculty of the University of Zurich. We gratefully acknowledge all the technical support from our lab mates who contributed to over the years to the development and maintenance of the DR and DIO-P colonies.

## Declaration of generative AI and AI-assisted technologies in the manuscript preparation process

Statement: During the preparation of this work the author(s) used Claude in order to correct for grammatic mistakes and style improvement. After using this tool, the authors reviewed and edited the content as needed and take full responsibility for the content of the published article.

## Notes

### Competing Interest Statement

The authors have declared no competing interest.

