## Supplemental Figure 1 for "Depletion of astrocyte inflammatory pathway in the arcuate nucleus of the hypothalamus is sufficient to prevent the diet-induced metabolic alterations of polygenically predisposed obese rats"

**Supplementary Figure 1**

**MBH astrocytic IKKβ depletion before HED exposure reduces food intake and alters meal patterning in ad libitum fed DIO-P rats.**
(**A-D**) Meal pattern analysis during *ad libitum* feeding: (**A**) cumulative food intake in g, (**B**) meal size in g, (**C**) meal duration in minutes and (**D**) meal number over the 12h light phase, 12h dark phase or the entire 24h period.
(**E-H**) Time course of energy expenditure during light-phase and dark-phase of *ad libitum* fed rats. (**E**) Measurement of EE (in kJ/h) in hourly bouts during the light and dark phase (grey area) and (**F**) Expressed as the sum of the light, dark or entire 24h period. EE was normalized to body composition (lean body mass+0.2 fat mass). (**G**) Measurement of RER in hourly bouts during the light and dark phase (grey area) and (**H**) expressed as the average of the light, dark or entire 24h period. Data are represented as mean ± SEM (n=6-8/group) and were analyzed using a 2-way (Factors: phenotype, AAV) or 3-way ANOVA (Factors: phenotype, AAV and time) followed by Tukey's post-hoc test. *$p<0.05$, **$p<0.01$, ***$p<0.001$, ****$p<0.001$.

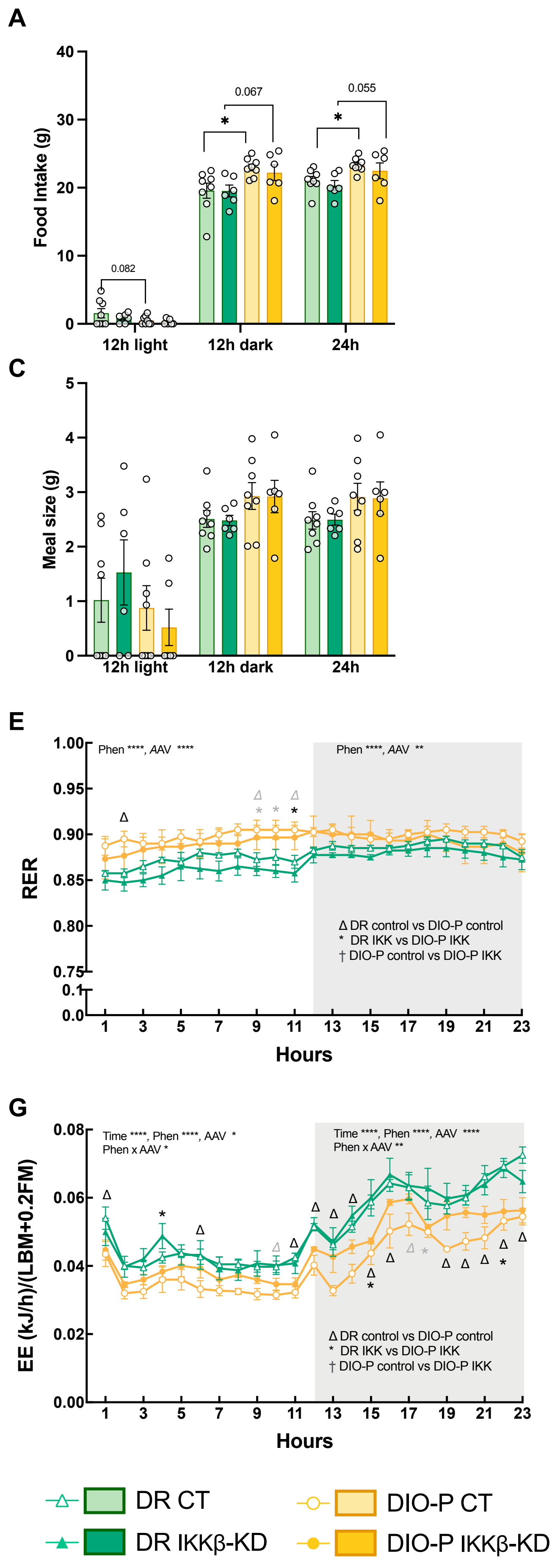
A
Food Intake (g)
0.082
0.067
0.055
12h light
12h dark
24h
B
Meal duration (min)
C
Meal size (g)
D
Meal Number
E
RER
Phen ****, AAV ****
Phen ****, AAV **
Δ DR control vs DIO-P control
* DR IKK vs DIO-P IKK
† DIO-P control vs DIO-P IKK
Hours
F
RER (avg/12h)
0.070
Light
Dark
24h
G
EE (kJ/h)/(LBM+0.2FM)
Time ****, Phen ****, AAV *
Phen x AAV *
Time ****, Phen ****, AAV ****
Phen x AAV **
Δ DR control vs DIO-P control
* DR IKK vs DIO-P IKK
† DIO-P control vs DIO-P IKK
Hours
H
EE (kJ/12h)/(LBM+0.2FM)
0.070
DR CT
DR IKKβ-KD
DIO-P CT
DIO-P IKKβ-KD
